# Mechanisms for Stable Bump Activity, Winner-Take-All and Group Selection in Neuronal Spiking Networks

**DOI:** 10.1101/077396

**Authors:** Yanqing Chen

## Abstract

A major function of central nervous systems is to discriminate different categories or types of sensory input. Neuronal networks accomplish such tasks by learning different sensory maps at several stages of neural hierarchy, such that different neurons fire selectively to reflect different internal or external patterns and states. The exact mechanisms of such map formation processes in the brain are not totally understood. Here we study the mechanism by which a simple recurrent/reentrant neuronal network accomplish group selection and discrimination to different inputs in order to generate sensory maps. We describe the conditions and mechanism of transition from a rhythmic epileptic state (in which all neurons fire synchronized and indiscriminately to any input) to a winner-take-all state in which only a subset of neurons fire for a specific input. We prove an analytic condition under which a stable bump solution and a soft winner-take-all state can emerge from the local recurrent excitation-inhibition interactions in a three-layer spiking network with distinct excitatory and inhibitory populations, and demonstrate the importance of surround inhibitory connection topology on the stability of dynamic patterns in spiking neural network.

## 1 Introduction

Facing with vast amount of multi-sensory information, Central Nervous System (CNS) seems to process only a small subset of those inputs at any given time, no matter whether they come from external or internal sources. How brain selectively processes such large number of inputs and maintains a unified perception remains a mystery. At the level of neuronal networks, a network in which all neurons respond the same to all stimuli would convey no information about the stimulus. In order to be useful, neurons must come to respond differentially to variety of incoming signals. Many neural models and theories have been proposed to account for such ability. Winner-Take-All (WTA) network is one of such proposed mechanisms for developing feature selectivity through competition in simple recurrent networks, and it has received much attention on both theoretical and experimental grounds. The primary theoretical justification is the ability of such networks to explain how the maps, which are ubiquitous in the cerebral cortex, can arise (Kohonen, 1982; Goodhill, 2007). WTA networks can also explain how a network can come to make useful distinctions between its inputs. WTA networks coupled with synaptic learning rules and homoestatic plasticity can explain how this takes place in a self-organized fashion from an initially undifferentiated state. Finally, WTA models are often employed at the behavioral level in theoretical models of higher-level cognitive phenomenon such as action-selection and attention (Itti & Koch, 2001; Walther & Koch, 2006).

Another mechanism proposed for feature selectivity is the phenomenon of spatially localized bumps in neuronal networks (Somers, Nelson, & Sur, 1995; Laing & Chow, 2001). If we view multiple neurons within a bump as mulitiple-winners of excitatory and inhibitory competition, bump activity in spiking networks can be treated as a soft WTA or k-Winner-Take-All phenomenon (see Maass, 2000 for their definition). In this paper we use

Winner-Take-All (WTA) and “bump activity” inter-changeably to describe the same stable group activity that arises from inter-connected excitatory-inhibitory neuronal networks. On a more general level, both bump activity and WTA phenomenon can be viewed as a type of pattern formation process in networks of excitatory and inhibitory neurons (for example, patterns of stable grid in Wilson & Cowan, 1973 and Ermentrout & Cowan, 1979), and an example of activity dependent neuronal group selection process (Edelman, 1987).

Population rate-based WTA models have been extensively studied and are well understood (Dayan & Abbott, 2001). But, the connections between rate models to the real biological neural systems are not direct, because they are different from the real nervous systems whose neurons are spiking. So it is necessary to study the networks of spiking neurons, such that the biological interpretation of spike models can be more directly linked to real nervous systems. Modeling and understanding spiking networks is not simple because spiking neurons are highly nonlinear and their action potentials are discrete. As a result, it is always more difficult to obtain analytical solutions for spiking firing properties than rate models.

Analysis has shown conductance-based spiking models can be approximated by simple rate models under certain conditions (such as in an asynchronous state in Shriki et al. Neural Computation, 2003). This approach has been applied to the study of hyper-column in a spiking model of visual cortex (Shriki et al., 2003). The orientation selectivity in their study, is modeled as the appearance of a unimodal “bump”-like spiking activity in a ring-connected spiking network, similar to an earlier study (Laing & Chow, 2001). Both approaches applied approximations from the rate models and used Fourier analysis to calculate the conditions for the appearances of bump activity. Recent work specifically studied recurrent spiking WTA networks, which are closer to real biological systems than previous rate models (Ref. Rutishauser & Douglas, 2009, 2011). Even though these newer network models can receive spike input and generate spike output, their network structures are still very simplified. For example, excitatory and inhibitory neurons are modeled into one single population (Laing & Chow, 2001), and inhibitory population are reduced into one unit (Rutishauser & Douglas, 2011), or removed altogether and modeled as direct inhibitory connections among excitatory neurons (Oster, Douglas, & Liu, 2009).

In a recent report we presented a robust and more biologically-realistic WTA network structure with distinct excitatory and inhibitory populations with arbitrary number of units (Chen, McKinstry, & Edelman, 2013). Each neuron type has very detailed biological parameters to model different neuronal transmitters and receptor types similar to previous work (Izhikevich & Edelman, 2008). We showed that surround inhibition and longer time constants from NMDA and GABAb conductances are sufficient to achieve stable “bump” spiking activity in a selected winner neuronal group while all the other neurons are inhibited and quiet. However, detailed biological properties, such as STSP (short-term synaptic plasticity), NMDA voltage gating etc., prevented a formal analytical analysis of the whole model. Also, it is not clear any of those biological details or a specific type of synaptic connections are crucial for the emergence of bump activity.

To identify the most important mechanistic factors for the spiking WTA networks, here we study a simplified spiking network after some biological details are removed. For example, based upon what we have noticed previously, turning off STSP, NMDA voltage-gating and excitatory-to-excitatory connections does not change the overall properties of WTA phenomenon. On the other hand, we preserve some important biological features such as the four different synaptic connections and conductance types (AMPA, NMDA, GABAa, GABAb), because we found that these four individual conductance types contribute to different aspects of the “bump” stability. By examining functions of these individual conductances and the topologies of excitatory-inhibitory connectivity, we provide a detailed analysis of the conditions on which a stable bump activity can emerge from this reentrant spiking network. Our analysis thus provide a mechanistic analysis on how a neuronal group selection process can occur in an activity dependent manner in neural systems.

## 2 Analysis of the Dynamic Firing Patterns in a Basic Neuronal Spiking Network

### 2.1 Network structure

Here we analyze a basic 3-layer spiking neuronal network with different neuron types with realistic biological parameters. The first layer of excitatory neurons (IN – input cells) takes input signals (e.g., arbitrary analog patterns) and translates them into spiking activity. The input signal we considere here in this paper is a type of unstructured random currents evenly distributed within a certain range and injected into the 100 input neurons (IN). IN cells are randomly connected to the next excitatory layer (E) with initial weights evenly distributed between 0 and a maximal value. The random input currents and random connections to the excitatory layer we analyzed here provide a baseline condition in which we test how the recurrent/reentrant connectivity between excitatory and inhibitory neurons by themselves can accomplish winner-take-all competition to random but unstructured input patterns (without obvious firing-rate differences among input neurons) and without synaptic modifications. The successful WTA network structure then can be trained to discriminate more complex and structured patterns through spike-timing dependent learning rules such as STDP. Such learning process will modify the synapses between these two excitatory types so that a selected E and I neurons (the WTA group) will respond to preferred input patterns more quickly for practical applications, which we have demonstrated in a previous report (Chen, McKinstry, & Edelman, 2013).

**Figure 1:**
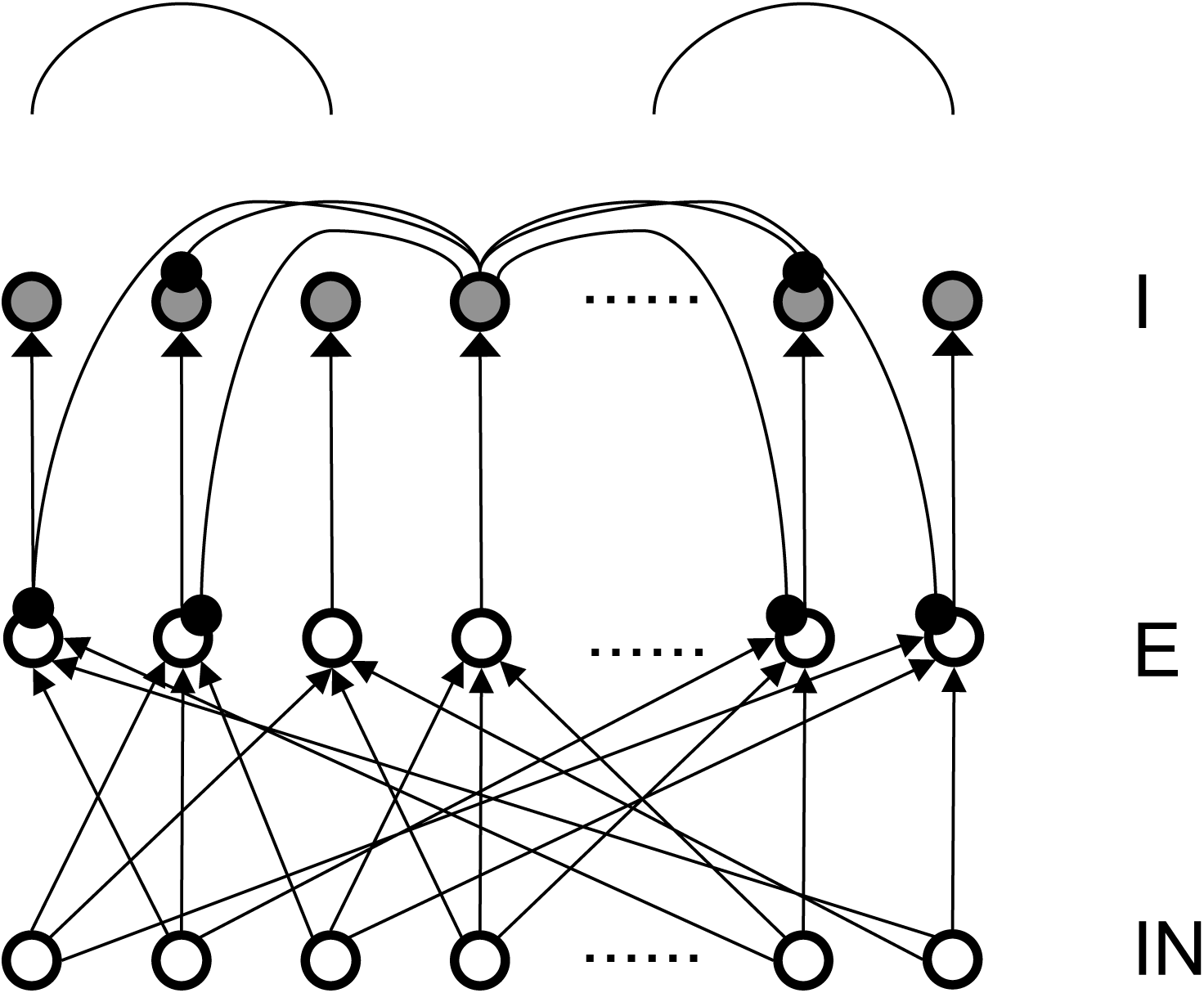
Structure of the basic 3-layer spiking network and a schematic plot of the “surround inhibition” connectivity that supports winner-take-all phenomenon. IN – thalamocortical input neurons, E – Excitatory pyramidal neurons, I – Inhibitory neurons. We chose 100 input (IN) cells, 400 E cells and 400 I cells for total of 900 neurons in the analysis model presented here. Input layer to excitatory layer (IN to E) are all-to-all random connected, excitatory to inhibitory layers are narrow and were simplified into one-to-one connection in our analysis. Inhibitory connections are surround type, that is, I cells do not inhibit its nearest neighboring I and E cells, but only distant surrounding neurons. This connectivity is implemented as two cosine peaks with a flat gap (zero value connectivity) in between. We call this specific network connectivity as surround inhibition type for the one dimensional case and it is a simplified version of the two-dimensional Central-Annual-Surround (CAS) type of topology we described before (Chen, McKinstry, & Edelman, 2013).

We also implemented the above network using adaptive exponential spiking models and obtained similar results. For simplicity the analysis below uses the Izhikevich model (Izhikevich & Edelman, 2008), and excitatory (E) and inhibitory (I) neurons use the same parameters in the following equation:

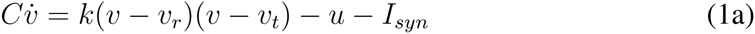

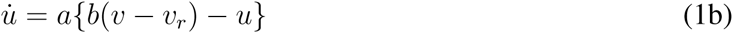

Parameters in these equations are the same as explained before (Izhikevich & Edelman, 2008). That is, *v* is the membrane voltage in millivolts (*mV*), *C* is the membrane capacitance, *v*_*r*_ is the neuron’s resting potential, *v*_*t*_ refers to its threshold potential, *u* represents the recovery variable defined as the difference of all inward and outward voltage-gate currents. *I*_*syn*_ is the synaptic current (in *pA*) originated from spike input from other neurons. *a* and *b* are different constants. When the membrane voltage reaches a threshold, i.e., *v > v*_*peak*_, the model is said to generate a spike, and two variables in Eq.1a and Eq.1b are reset according to *v ← c* and *u ← u* + *d* while *c* and *d* are parameters for different cell type.

We use a simplified synaptic current form with four basic conductances from *AMPA*, *NMDA*, *GABA*_*A*_ and *GABA*_*B*_ channels. For simplification, voltage-gating of NMDA channel is reduced to a constant factor. This is done through calculating an average number for the voltage-gating term for the NMDA conductance (i.e., [(*v* + 80)*/*60]^2^*/*[1 + ((*v* + 80)*/*60)^2^] on Page 11 of the Supplementary Information in Izhikevich & Edelman (2008)) for the normal range of voltages: *v* = [*−*60, 60]), and the result is equivalent to a voltage-independent NMDA channel with smaller gain factor than AMPA channels (see Appendix for the individual conductance gain factors we used). So synaptic current *I*_*syn*_ is composed of four different current types originated from those four conductances multiplied with the voltage differences between their individual reversal potentials:

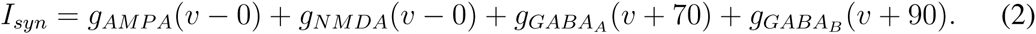

Shown in the above equation, reversal potentials of *AMPA* and *NMDA* channels are 0 and reversal potentials for *GABA*_*A*_ and *GABA*_*B*_ channels are *−*70*mV* and *−*90*mV* respectively.

As described before, each conductance has exponential decay with different time constants in millisecond (ms):

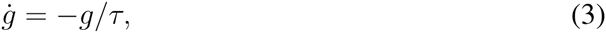

while *τ* = 5, 150, 6, and 150 for the *AMPA*, *NMDA*, *GABA*_*A*_ and *GABA*_*B*_ channels respectively.

To simplify the analysis, there are equal numbers (400 in all the subsequent analysis) of excitatory (E) and inhibitory (I) neurons in our basic network model in Figure 1, although their numbers can be in any ratio. We also explored different types of connection topologies in the connections from excitatory to inhibitory neurons (E to I), the reentrant inhibition from basket cells to pyramidal neurons (I to E) and the inhibitory connections within basket cells themselves (I to I). In our study, Inhibitory to Excitatory and Inhibitory to Inhibitory connections are kept the same topological type and total weights are kept equal. Throughout the simulation the total connection weights to each neuron are normalized to be a constant for each connection type. The total weights for each connection type (E to I and I to E) are two parameters we explored systematically. As a first step, we firstly only consider one type of inhibitory conductance (*GABA*_*A*_) to obtain analytical solutions for the conditions of Winner-Take-All state. *GABA*_*B*_ conductances are added after an analytical solution is found, a comparison of the transition plots can be found in the Appendix.

To classify different types of spike dynamics for the surround inhibition network in Figure 1, for each neuron, we record the number of spikes between 2 and 3 seconds after the simulation had reached steady state without synaptic plasticity (STDP off). We then characterize the behavior of the network by the maximum number of spikes generated by any excitatory neuron. Figure 2 (e) plots this maximal firing rate for every combination of the E to I weights versus the I to E weights. The analysis is repeated and plotted in Figure 2(f) for the inhibitory population.

Figure 2 shows different types of dynamic firing patterns in the 2-dimensional parameter space. When only one type of connection weight (excitatory or inhibitory) is high but the other weight is low, either excitatory or inhibitory neurons are in a quasi-random / rhythmic state in which one group of neurons fires in high Gamma frequency range (*>* 40Hz, see Figure 2 (a) and (g)). When both connection weights are relatively high (see Fig.2(c)), both excitatory and inhibitory neurons have high maximal firing rates where excitatory neurons have a maximal firing rate larger than 35Hz and inhibitory neurons have a maximal firing rate of larger than 100Hz. If we look at the corresponding spike raster plot in Fig2 (c), only a subset of excitatory and inhibitory neurons maintain such high firing rates while majority of other neurons are silent. We call this Winner-Take-All (WTA) state in which only a small subset of neuronal groups persistently fire high frequency and keep the rest of neurons from firing using surround inhibition. The region of the parameter space with such WTA states is delineated by the right curve in Fig.2(d) where the maximal firing rate of excitatory neurons increased to greater than 35Hz from lower firing rates in the middle region (from the blue area in Fig. 2(e) transition to the red area on the top right), and roughly corresponds to a similar increase of maximal firing rate to above 100Hz for inhibitory neurons in Fig.2 (f).

**Figure 2:**
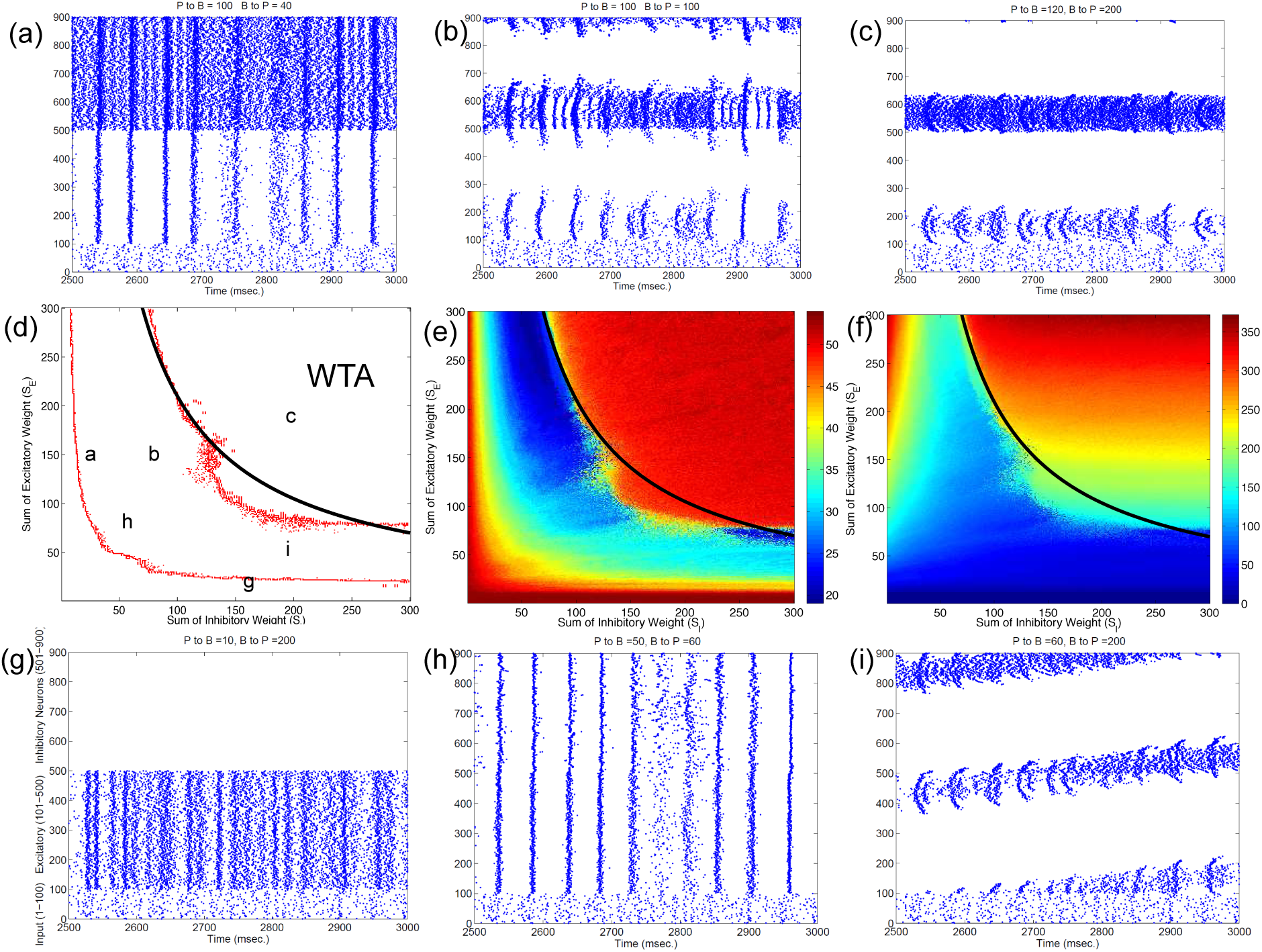
Classification of different dynamic spiking patterns in the surround inhibition network we defined in Figure 1 and the phase diagram for the transition between different firing states.(a) to (c) and (g) to (i) are raster plots which show all spikes within a half second interval for each neuron in the network. (e) and (f) are maximal firing rates of excitatory and inhibitory neurons in the 2-d parameter space (total excitatory weight in y-axis and total inhibitory weight in the x-axis). Red points in (d) are transition curves constructed from (e) where maximal firing rate of excitatory neurons changed from below to above 40Hz. (a) to (c) and (g) to (i) are example spiking patterns under their specific parameter combinations which are marked on the 2-d plot in (d). Subplot (c) represents a winner-take-all (WTA) state where only a small group of excitatory and its corresponding inhibitory neurons fire persistently while others are silent. Black curves on the middle row – (d), (e) and (f) are the same analytic transition condition for the WTA region based upon the analysis described in the Appendix (Eq. (19)). Basically it is a curve where *S_E_ · S*_*I*_ equals to a constant defined by Eq. (19), and it delineates the Winner-Take-All region (i.e., max firing rate large than 40Hz for only a selected group of excitatory neurons) in the parameter space very well for both the excitatory and inhibitory neurons.

Subplots (a), (b), (h) and (i) in Fig. 2 all belong to an intermediate region in the parameter space in Fig.2(d) between two curves where maximal firing rates for both excitatory and inhibitory neurons are relatively low. Within this parameter range, excitatory and inhibitory neurons are either quasi-synchronized (Fig.2(a)) or precisely synchronized and firing rhythmically (Fig.2(h)), or exhibit moving bump activity (Fig.2(i)) or as combinations of rhythmic and moving bump activity. In all these cases, single excitatory neuron cannot maintain a stable high gamma frequency spiking activity unless connection weights are changed, moving to the WTA region on the top-right of the second curve in Fig.2(d). Fig. 2(d) thus provides a phase diagram for the neuronal network defined in Fig.1.

Notice that this maximal firing rate is not the neuron’s instantaneous firing rate, but is the total number of spikes within a 1 second window. This definition is useful to discriminate a stable high firing rate neuron versus a neuron firing a short burst less than 1 second and then becoming quiet (especially for stable vs. traveling activity, see Figure 2(c) vs. (i)).

Figure 3 summarizes patterns of spike dynamics with different connection topologies. Compared to the surround inhibition type analyzed above, all the other connection types do not support a Winner-Take-All state manifested as stable bump activity shown in Fig.2(c). This is because under those connection types, excitatory and inhibitory neurons cannot maintain high maximal firing rates when both excitatory and inhibitory weights are high and did not have a red area on the upper-right region shown in Fig.2 (e) and (f). The most common firing patterns for those connection types are quasi-rhythmic firings in the 10 to 20 Hz range for excitatory neurons resembling an epileptic state while some short burst of unstable bump activity in inhibitory neurons. Our results suggest that, among different types of connectivity topologies we analyzed, only surround inhibition can generate a stable bump spiking activity and maintain a WTA state.

**Figure 3:**
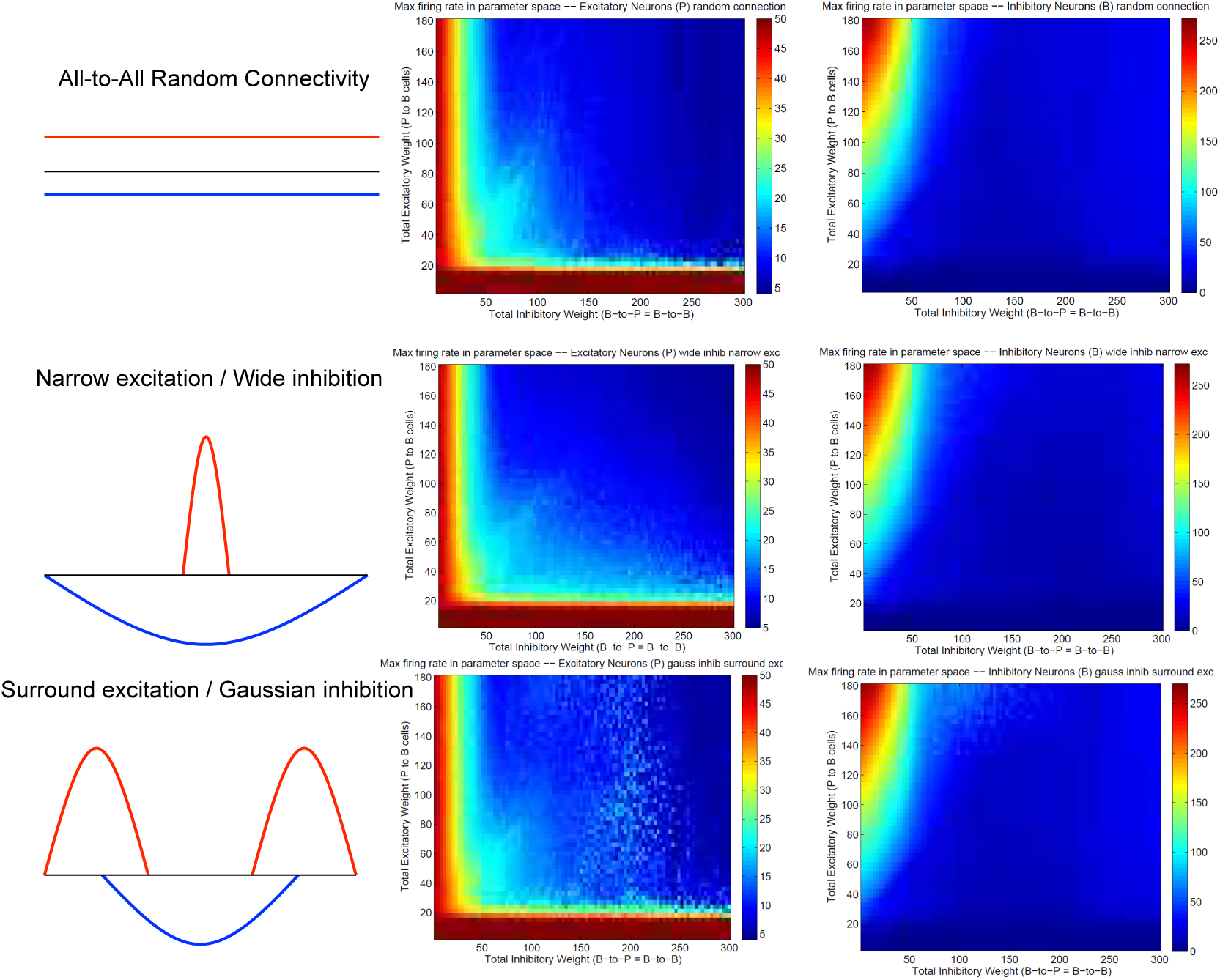
Spiking dynamics under different connection types other than the surround inhibition type defined in Figure 1. Each row represents a connectivity type and the middle and right columns are maximal firing rate of excitatory and inhibitory neurons under specific connectivity. These plots were calculated the same way as Figure 2(e) and (f). Notice that all three connectivity types here do not support a winner-take-all (WTA) region in the parameter space (no red region in the upper-right corner). It exists in Figure 2 (e) and (f) as a red region representing a high individual maximal firing rate state when both excitatory and inhibitory weights are relatively high, but it is always absent here on the top right of the 2-d parameter space.

## 3 Mechanism of Winner-Take-All neuronal group selection and emergence of bump activity

Our above analysis suggests that surround inhibitory topology supports emergence of bump activity. To explore the mechanism of WTA and which neuronal properties are essential for such behavior, we applied the same analysis as in Fig. 2 to neuronal network in Fig.1 when Short-Term-Synaptic-Plasticity (STSP) or NMDA voltage-gating is on, or change excitatory and inhibitory neurons’ parameters to different type. In all cases, a similar WTA region was found for every conditions, even though the transition curves that delineate the emergence of stable bump activity are shifted to different positions in the parameter space. We also analyzed the same neuronal network with a different set of individual spiking models, i.e., the adaptive exponential models and found the similar WTA region as long as the topology of the inhibitory connections are surround type. These analyses suggest that detailed neuronal properties such as exact models of the spiking neuron, STSP or NMDA gating etc., are likely not fundamental for the existence of stable bump activity, but the type of connectivity topology (i.e., surround inhibition) is more important for such behavior.

Both the Izhikevich neural model and the adaptive exponential model we used are conductance based with models of inhibitory and excitatory currents of different time scales. So we suspect that different time constants of NMDA, AMPA, GABAa and GABAb channel conductance might play some role for the emergence of bump activity. To demonstrate this, Figure 4 shows the time evolutions of AMPA, NMDA and GABAa currents along with the spiking activity in the simplified network in Figure 1 starting from a zero conductance initial condition. It demonstrates the detailed transition from a rhythmic synchronized firing state into a stable bump activity. Looking at detailed dynamic changes of the individual excitatory and inhibitory currents should shed light on how the transition is occurred.

**Figure 4:**
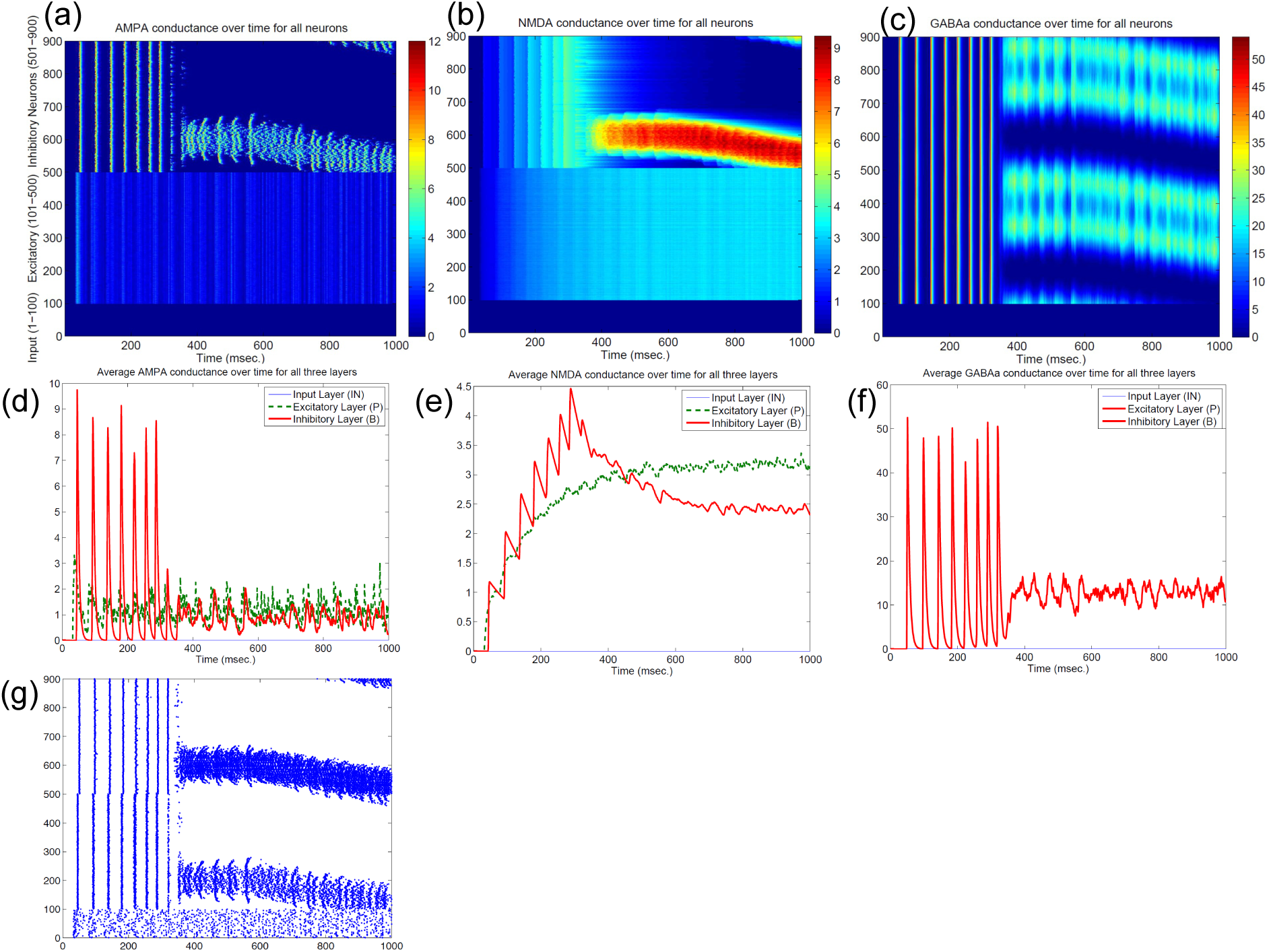
Transition from a synchronized rhythmic firing state to a winner-take-all bump spiking activity (g). In this simplified surround inhibition network, short-term synaptic plasticity (STSP) and NMDA gating effects are removed, GABAb conductance is set to be zero and individual excitatory and inhibitory spiking models are changed to have the same biological parameter set. First row shows the 1-second spatial-temporal evolutions of individual conductance for AMPA – (a), NMDA – (b) and GABAa (c). Second row show the averaged conductance over time for excitatory and inhibitory cells separately. Notice that bump activity in AMPA and NMDA conductance appear in inhibitory neurons first (see (a) and (b)), while both conductances are spatially uniform even after spiking bump activity emerged after about 400 msec. The fact that spatial uniformity is destroyed in GABAa conductance first in (c) suggests that inhibitory neurons might show transition into winner-take-all state early and then bias the transition in excitatory cells.

From Figure 4 we can see, differences in time constants will determine how fast a specific channel conductance returns back to zero after a burst of spiking activity. For excitatory neurons specifically, because of its short time constants (6 msec.), AMPA current fluctuates around a similar level with large variances. NMDA currents, on the other hand, are accumulating to higher levels because of longer time constant (150 msec.) even though both currents are generated by the same spiking input from the input neurons. Similar phenomenon can be seen for inhibitory neurons. When those neurons fire rhythmically before about 300 millisecond, AMPA conductance jumps to high level (from 7 to 9 nS) after each spike then drops down to zero very fast (red curve in Fig.4(d)), while NMDA conductance only drops a small amount each cycle and overall level still increases to much higher value (red curve in Fig. 4(e)). Initially inhibitory neurons fire after excitatory neurons in each rhythmic cycle and they synchronize to each other with a time delay. If excitatory to inhibitory weights (E to I) are larger than a certain value, such that NMDA currents for inhibitory neurons increase faster than excitatory neurons (Fig. 4(e)), the delay between inhibitory and excitatory neurons diminishes and GABAa currents become effective within the same cycle to inhibit other neurons. As a result some inhibitory and excitatory neurons stop firing in the rhythmic cycle, eventually lead to a winner group that persistently fires and shuts off their surrounding neighbors. Notice that in the simplified model in Figure 4, GABAb (with longer time constant of 150 ms.) currents are omitted and set to zero, which lead to a moving bump activity for this specific parameter set. If GABAb conductance is restored to the original level as in the full model, bump activity becomes stable. It implies that time constant of GABAa and GABAb channel conductance is related to the stability of the bump activity,

As a summary, we think the combinations of long and short time constants from excitatory and inhibitory conductance plus the surround inhibitory connectivity support a mechanism for emergence of bump activity and winner-take-all phenomenon in this basic spiking neuronal network. This neuronal group selection mechanism provides a basis for modeling learning and map-formation process for sensory motor integration and other higher cognitive processes.

## 4 Analytical Analysis of the Transition Curve for WTA phenomenon

### 4.1 Differentiation in Inhibitory Conductances Lead to Spiking Activity Differentiation and Neuronal Group Selection

To identify the mechanism about the emergence of bump activity in spiking network, based upon the transition plots shown in Figure 4, we look at voltages, conductances of all 400 excitatory neurons at one specific time point (t=980 msec. in Figure 4). Figure 5 shows AMPA, NMDA, GABAa conductances long with voltages for all those excitatory neurons at this specific time point. From the network structure defined in Figure 1, we know that the excitatory conductances (AMPA and NMDA) are determined by excitatory synapses from the input layer (because we omitted self-excitation), where inhibitory conductances (GABAa and GABAb) are only determined (triggered) by spikes from inhibitory neurons onto those excitatory neurons. Because of the uniform random connection from input layer, AMPA and NMDA conductances are around the same level and undifferentiated for all excitatory neurons. From Figure 5, voltages are above threshold and neurons fire only at locations where GABAa conductances below a certain value. So in order for a bump to emerge and a subgroup of neurons selected to be active, GABAa or GABAb conductances have to be differentiated, i.e., they have to be small for some neurons and remain high for all other neurons. We suspect this condition can only be met by surround type of inhibitory connectivity. It is obvious that lowest GABA conductances lead to highest firing rate for excitatory neurons. Since we have local feedforward excitation to inhibitory neurons in our network, the bump area in inhibitory neurons with highest firing rate should also have lowest inhibitory conductances. This difference in GABA conductances is true for both excitatory and inhibitory neurons because GABA conductances are determined by the same inhibitory spikes. This suggests that in order for the bump to emerge, local inhibition to the nearest neighbors should be lower than inhibition to neurons outside of bump. Notice the three other inhibition topology in Figure 3 all have peak (or flat) inhibition locally, so even if a neuronal group emerge with firing highest firing rate, the strong local inhibition will force their firing rate to decrease, and let the other sub-threshold neurons to fire. So this is likely the reason why we did not obtain stable bump activity using those inhibition connectivities. On the other hand, surround inhibition type defined in Figure 1 might be the most simple form of inhibition topology that could let a bump emerge and stabilize. Below analysis will further prove this point.

**Figure 5:**
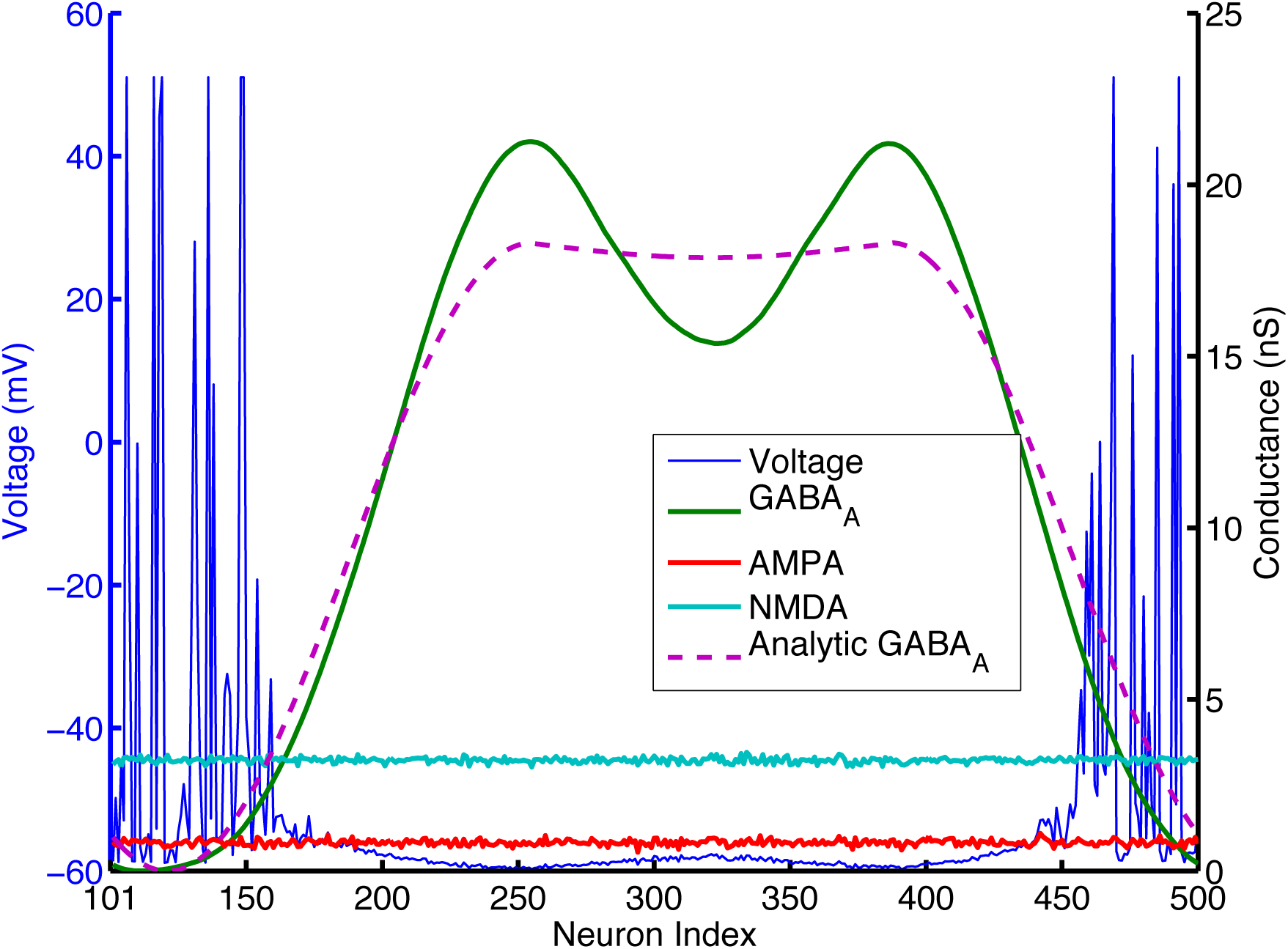
*AMPA, NMDA, GABA*_*A*_ conductances and voltages for all excitatory neurons (cell index 101 to 500 in Fig. 1) at one specific time point (t=980 msec. in Figure 4). AMPA and NMDA conductances are basically flat across all excitatory neurons here, while *GABA*_*A*_ conductances are close to 0 for neurons around number 101 and 500, and reach high values else where. Pink dashed lines are analytic GABA conductances from Eq. (10) in the Appendix and derived from surround inhibition topology, and is a good match for the actual numerical simulation. Notice that excitatory neurons fire spikes and have above threshold voltage values (blue lines) only within neighborhood of neurons having *GABA*_*A*_ conductances close to 0 and below a certain value.

Comparing with models using negative weights to represent inhibitory connections, a major difference in our more biologically realistic spiking models is that synaptic weights and excitatory/inhibitory conductances are all positive. From Eq. (2) we can see, it is only because of the differences in reversal potentials between excitatory and inhibitory channels, the current generated by excitatory and inhibitory conductances could have different signs (excitatory current coming into the neuron and inhibitory current coming out of neuron). In order for a neuron to fire, synaptic current has to be below a threshold *I_syn_ < −I*_*th*_ where *I*_*th*_ is about 100pA (*I*_*syn*_ has to be negative for a neuron to fire because it was defined as an outward current in Eq (1a)). Firing threshold *I*_*th*_ can be found from calculating F-I curve for the specific spiking neuron model we used. Figure 9 (in the Appendix) plots the firing rates versus amount of injected current (equivalent to *−I*_*syn*_) for the Izhikevich neuron model result from numerical simulations. It shows that neurons start to fire when absolute value of the injected current is above 100pA and then increase their firing rate approximately linearly until above 100Hz. we can use this information to simplify the spiking activity into a rate model. As indicated on the last paragraph, AMPA and NMDA conductances are approximately uniform for excitatory neurons and they can not contribute to the differences in firing rates, so in order for the excitatory population to fire differentially, the difference between highest and lowest GABA conductances for individual neurons has to be larger than a certain value. This value can be estimated using Eq (2). If *min*(*g*_*GABAA*_) is 0, for a resting potential of *v*_*r*_ = *−*60*mV*, *GABA*_*A*_ conductance has to be larger than the following value so injected synaptic current *−I*_*syn*_ will be below the firing threshold *I*_*th*_:

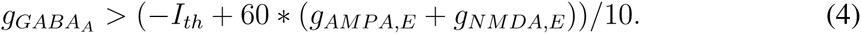

In Figure 5, *g*_*AMP A,E*_ +*g*_*NMDA,E*_ for excitatory neurons is around 4nS (Appendix will show how this value can be estimated analytically), so *g*_*GABAA*_ has to be larger than 14nS to keep sub-threshold neurons from firing. This number is consistent with the result plotted on Figure 5 and Figure 4 that neurons fire and form a bump area where GABA conductances are below 14nS and areas with GABA conductance larger than 14nS are completely quiet.

If we consider both the *GABA*_*A*_ and *GABA*_*B*_ conductances based upon Eq. (2) and using the same idea as above, conditions for inhibitory conductances will be the following for the winner-take-all state:

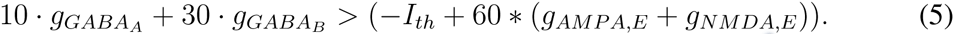

Eq. (4) and Eq. (5) can be used further to identify the exact condition for the WTA state and to locate the transition curve in Figure 2. Using two cosine bumps as surround inhibitory connection weights, Appendix gives the analytic form of GABA distribution for a bump solution for neurons connected one dimensionally and uses it to obtain analytic conditions for the WTA state in the parameter space (see Eq. (19) and Eq. (20)). Such analytic conditions are expressed as formulas combining single neuron property and the conductance parameters (such as time constants, gain factors for different inhibitory and excitatory conductances). Based upon these formulas we can locate the Winner-Take-All and bump activity in the parameter space fairly precisely (see black curves in Fig. 2 and the white curve in Fig. 10 in the Appendix), thus provide a mechanistic explanation for the emergence of winner-take-all state and stationary bump activity in this 3-layer spiking network we analyzed here.

### 4.2 Origin of Traveling Wave and Instability of Bump Activity – driven by AMPA conductances

Parameters in Figure 4 are located very near the transition curve in the parameter space (see Figure 2), so the bump is not spatially stable and moves across different neurons. To identify the origin of such instability, we selectively set AMPA gain of excitatory or inhibitory neurons to 0 in order to see their effects on the bump stability. This is equivalent to selectively block AMPA conductances in either excitatory or inhibitory neurons in real biological neural systems. We found that if AMPA conductances in inhibitory neurons are set to 0 but not in excitatory neurons, bumps become more or less stable. On the other hand, when AMPA conductances are blocked and set to 0 in excitatory neurons but not in inhibitory neurons, we can have a moving bump with a constant spatial speed (see Figure 6). In fact, we can estimate the moving speed of bumps based upon the parameters we defined in Figure 2. So we believe the source of the bump instability is from the AMPA conductances in inhibitory neurons.

**Figure 6:**
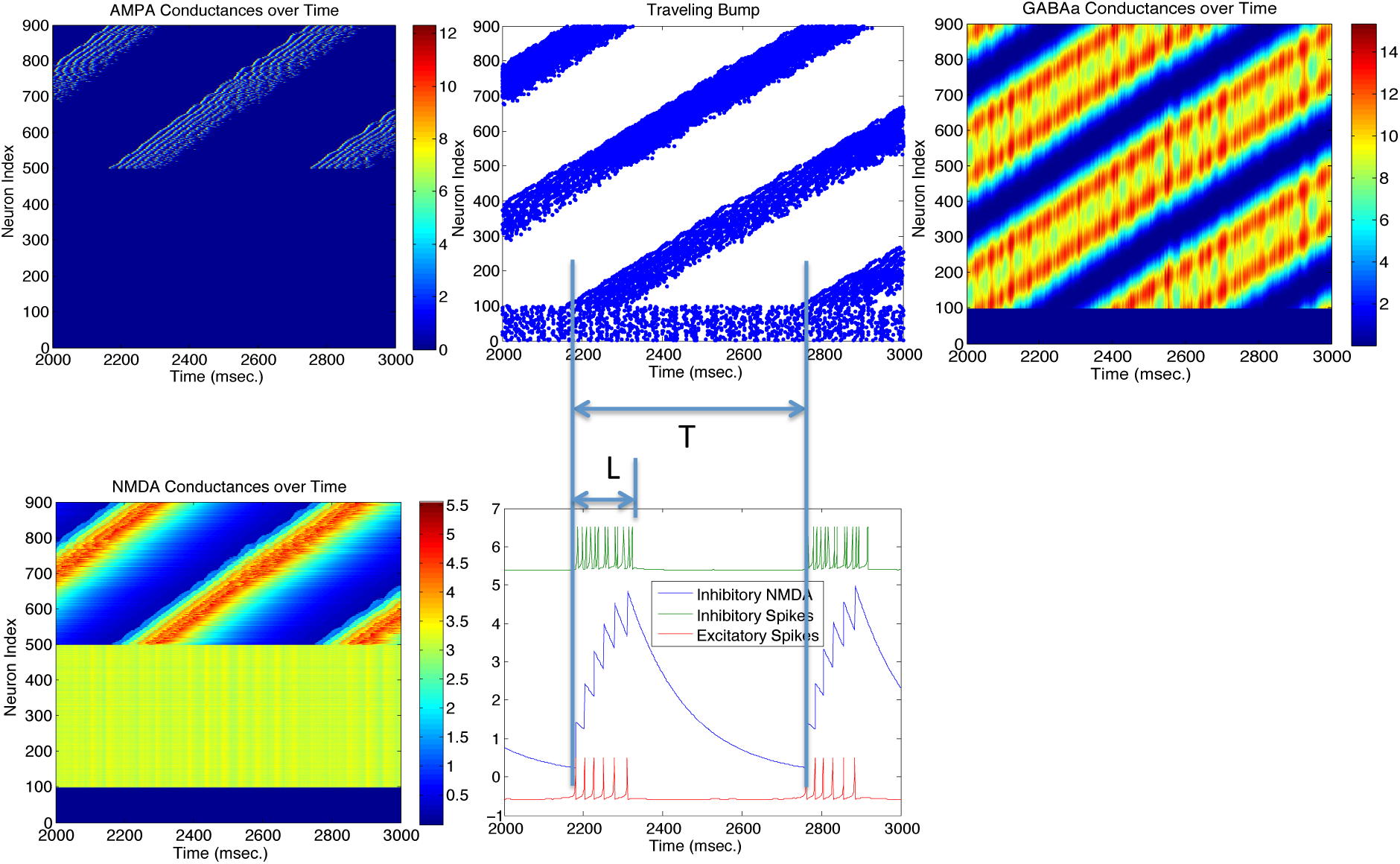
Stability of the bump activity is determined by AMPA conductances. Figures above show that spiking bump activity moves with a constant speed when AMPA conductances in excitatory neurons are blocked and set to 0 (Top left). The speed of the traveling “bump”, proportional to the length of firing interval for each neuron (L) and the period of the traveling wave (T), seems relate to the frequency of the inhibitory neurons (hence their AMPA conductances) and the delay time for the NMDA conductances when they reach maximal firing rate. When AMPA conductances are set to 0 in inhibitory neurons, bump activity stops moving and *T → ∞*.

## 5 Conclusion and Discussion

In this paper we derive global properties of spiking neuronal networks related to bump activity and Winner-Take-All state mainly through analysis of the dynamics of excitatory and inhibitory conductances. To achieve the analysis of collective behavior, individual spiking properties are approximated by its firing rate property such as the conductance/firing rate curve (Fig. (9) or F-I curves. In this regard, detailed properties of individual spiking model might not be crucial for global activity such as the emergence of bump and WTA state. For example, if we use different parameter sets for excitatory and inhibitory neurons such as change the inhibitory neurons to be basket cell type, we can still found the WTA region in the parameter space in Figure 2(d), but the exact location of the transition curve is shifted to a different place because basket neurons have different conductance/firing rate curve and different *I_th_, g_th_, k* values in Eq. (19). This could explain the transition curve in our full model with more detailed biological properties has the same functional form, but also in different location in the parameter space because it included more detailed single neuronal spiking properties such as NMDA voltage gating and STSP etc. In fact, we used adaptive exponential spiking model to substitute the Izhikevich neuron models and obtained similar phase plot and transition curve for the bump activity and the WTA state.

We suggest that all conductance-based spiking models with distinct excitatory and inhibitory populations could have the similar collective Winner-Take-All behavior as analyzed here. Detailed spiking model properties such as F-I curve and firing threshold (*I*_*th*_ and *g*_*th*_) would determine the exact location of the transition curve in Figure 2. Global connectivity topology and different time constants (dynamics) of excitatory and inhibitory conductances are likely to be the determinant of system-wide spiking activity patterns.

### 5.1 Importance of the Inhibitory Topology

The most important feature of our winner-take-all network is its surround inhibition topology. We chose two sine function peaks as surround inhibitory connection weights is just because of its mathematical convenience, since convolutions of sine/cosine functions are much easier to solve than other types of functions. In fact, we actually used Gaussian peaks or torus (for two-dimensional neuronal arrays) in our previous model (Chen et al. 2013) and obtained similar stable WTA results. We believe using other type of function for inhibitory connection would also work, as long as there is a low inhibitory weight locally. Comparing four different connection topologies from Fig. 2 and Fig. 3, the reason why only surround-inhibition supports stable bump activities is because its maximal inhibitory connection weight is not to the nearest neighbors, but to slightly distant neurons. This gap of zero inhibitory weight could be very small (e.g., *w* down to 0 and equivalent to a no-self-inhibition case), and we could still find solutions for stable bump or bumps (see Fig. (7) and Fig. (8)). So as long as there is a local valley of inhibitory weight, stationary bumps could emerge because only decreasing inhibition could allow a bump to sustain.

Mechanistically it appears that the most important requirement for a bump solution is the stable differentiation in inhibitory (*GABA*_*A*_ or *GABA*_*B*_) conductance distributions across the neuronal population. That is, for some neuronal groups, GABA conductances should be low to allow bumps to emerge and for the other neurons, they need to be high enough to keep the rest of neuronal population from firing spikes. As long as this condition is met, more detailed biological properties such as local self-excitation, short-term synaptic plasticity (STSP), voltage-gating of NMDA channels etc. can be added to the model without destroying the overall bump stability.

### 5.2 Why Traditional Center-Surround Topology might not Lead to Stable Bumps in Models with Distinct Excitatory and Inhibitory Populations

Previous rate-based population models (Dayan & Abbot, 2001) had most often used center-surround type of connection topology as shown in the middle row of Figure 3 (Narrow excitation / Wide inhibition), similarly many spiking models with excitatory / inhibitory conductances on the same units used the same topology (Laing & Chow, 2001). By virtue of the direct mathematical operations, narrow excitation and wide inhibition can lead to a “Mexican Hat” type of effective connectivity which supports winner-take-all in previous firing rate models. As we see from the analysis above, in biologically more realistic spiking models with distinct excitatory and inhibitory neuronal populations, multiple types of conductances cannot cancel each other easily because they are generated by precise spike timing and have different time constants. The “classical” center-surround topology can not guarantee a stable “Mexican Hat” type of net connectivity because of sensitive spike timing differences between different neurons. In fact, as shown in Figure 4, the emergence of winner-take-all in spiking networks is a direct result of precise spike-timing - the coincide of excitatory and inhibitory population firing spikes lead to a sub-population of inhibitory neurons fire earlier than the rest of populations which then let them suppress and shut off the other neurons in the network (see Figure 4g). This is the reason that we believe a surround-type of inhibitory topology is essential for a stable spiking WTA network because it can support the emergence of a winner-group without shutting themselves off too early.

In summary, WTA network analyzed here demonstrates how variability and randomness in spiking time of individual neurons can lead to global pattern changes and phase transition in collective neuronal groups. Analytic solutions for the phase transition curve provided in this paper will help to increase our understanding of the different functional roles of excitatory and inhibitory neural connections on the emergence and stability of firing patterns in the brain.

## Acknowledgments

Research supported by the Neurosciences Research Foundation.

## 6 Appendix

In this Appendix we present some detailed mathematical analyses of the surround inhibition network we defined in Figure 1, including an analytic approximation of the excitatory and inhibitory conductance distribution within the network, especially the GABA conductances. Based upon those results, we can identify the function (shape) and the exact location of the transition curve we found in Figure 2(d) which delineates the emergence of the stable bump activity and WTA state within the parameter space.

In the main text we suggest that surround inhibitory connectivity can support the emergence of bump activity. This is because this type of inhibitory topology can result in a non-uniform GABA distributions with stable and separated peak and valley in inhibitory conductances, and the difference in GABA peak and valley has to be larger than a certain value to selectively activate some groups of neuron (winning bump) while keeping the rest of neurons from firing (see Eq. (4)). Here we derive the exact functional form of the GABA conductance distribution based upon one specific surround inhibitory topology we used. From the functional forms we can identify what parameter combinations will satisfy Eq. (4), thus locate the transition curve in the parameter space.

Because we have wrap-around connections between neuronal populations, we use a continuous angular coordinate (from *−π* to *π*) to represent the spatial location of the excitatory and inhibitory neurons. This continuous coordinate can be viewed as the limit of infinite number of neurons. In the case of limited number of neurons (400 in each type in our examples here) as depicted in Figure 2, excitatory neurons number 101 to 500 can be viewed as located evenly from *−π* to *π* in this angular coordinate, same as the inhibitory neurons number 501 to 900.

From Eq. (3), we know that all conductances have exponential decay with their specific time constants. The final conductance distributions over time, however, are the result of self decay plus the convolution of the synaptic weight function and the input spikes multiplied by a gain-factor *G*:

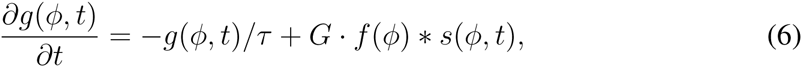

where *g*(*ϕ, t*) is a specific conductance at time *t* for neuron at location *ϕ*, and *s*(*ϕ, t*) are incoming spikes to this individual neuron represented by delta function, and *G* is a gain-factor equal to the amount of a specific conductance generated by one spike. If we consider excitatory conductances for excitatory neurons, because each excitatory neuron receives uniformly random input from all neurons from the input layer (see Figure 1), *f*_*IN*_ (*ϕ*) *∗ s*(*ϕ, t*) can be approximated by a constant, i.e., *f*_*IN*_ (*ϕ*) *∗ s*(*ϕ, t*) *≈ r_IN_ S*_*IN*_, where *r*_*IN*_ is the average firing rate of neurons in the input layer and *S*_*IN*_ is a pre-specified total synaptic weight from the input layer onto each excitatory neuron. Under this condition, Eq. (6) can be solved to identify the average excitatory conductances for excitatory neurons:

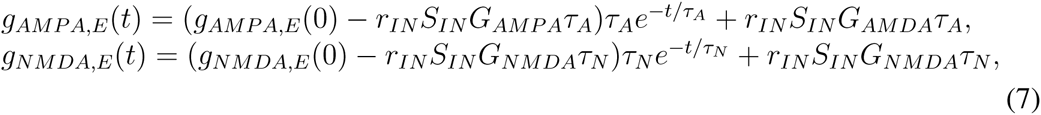

where *τ*_*A*_ = 5*ms.* and *τ*_*N*_ = 150*ms.* are time constants for AMPA and NMDA conductances respectively, and *G*_*AMP A*_ and *G*_*NMDA*_ are gain-factors representing the amount of AMPA and NMDA conductances generated by one spike. *g*_*AMP A,E*_ and *g*_*NMDA,E*_ are no longer functions of *ϕ* (neuron location or indices) because excitatory neurons receive uniformly random weights from input layer thus their excitatory conductances are spatially uniform and undifferentiated (see Figure 4(a) and (b) for neuron number 101 to 500). Comparing green curves in Figure 4(d) and (e), Eqs. (7) fit the time changes of average excitatory AMPA and NMDA conductances very well for the initial conditions we used (*g*_*AMP A,E*_ (0) = *g*_*NMDA,E*_ (0) = 0). Their steady-state values can be approximated by the last two terms in Eqs. (7), i.e.:

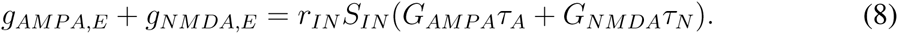

The total synaptic weight from input layer to each excitatory neuron we used is 100nS (*S*_*IN*_), *G*_*AMP A*_ is 0.15*nS/spike* and *G*_*NMDA*_ is 1/10 of that, for an average input frequency of about 14Hz (*r*_*IN*_), we can derive that the steady-state AMPA conductance is at 1nS and NMDA conductance is 3 times of that at 3nS. These values fit perfectly with the numerical results on Figure 5.

Next we derive the functional form for the inhibitory GABA conductances. The exact inhibitory connection topology we used has two cosine/sine connection weights with a gap in between (see top schema in Figure 6). This gap (size of 2*w*) is a parameter that can be adjusted and reflect a local region in which the originating neuron that sending out inhibitory synapses (located at 0 coordinate) does not inhibit. The neuron at location 0 sends inhibitory weights to the rest of neurons following the below function:

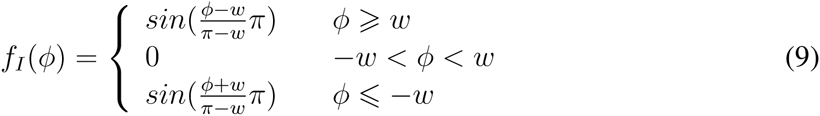

For a bump solution, we assume inhibitory neurons will fire within the gap region between *−w* and *w* with a firing rate of *r*_*I*_. This approximation is based upon the observation in Figure 5 that action potentials only occur when GABA conductances are below certain value and such condition could most likely to be satisfied within the region where the inhibitory weight is zero. Using firing rate approximation, GABA distribution is the convolution of the spikes between *−w* to *w* with function *f*_*I*_ (*ϕ*):

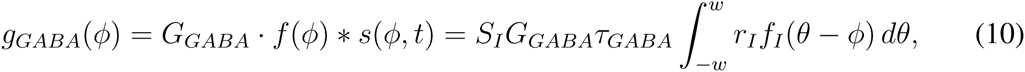

where *S*_*I*_ is the total synaptic inhibitory weight received by each neuron (excitatory or inhibitory) and is a parameter we pre-specify, *G*_*GABA*_ is the gain factor specify the amount of *GABA* conductance generated by one spike, and *τ*_*GABA*_ is the time constant and is 6*msec.* for *GABA*_*A*_ and 150*msec.* for *GABA*_*B*_ conductances respectively. These three terms are simplified to a constant in the following equations (set *C* = *r_I_S_I_G_GABA_τ*_*GABA*_).

**Figure 7:**
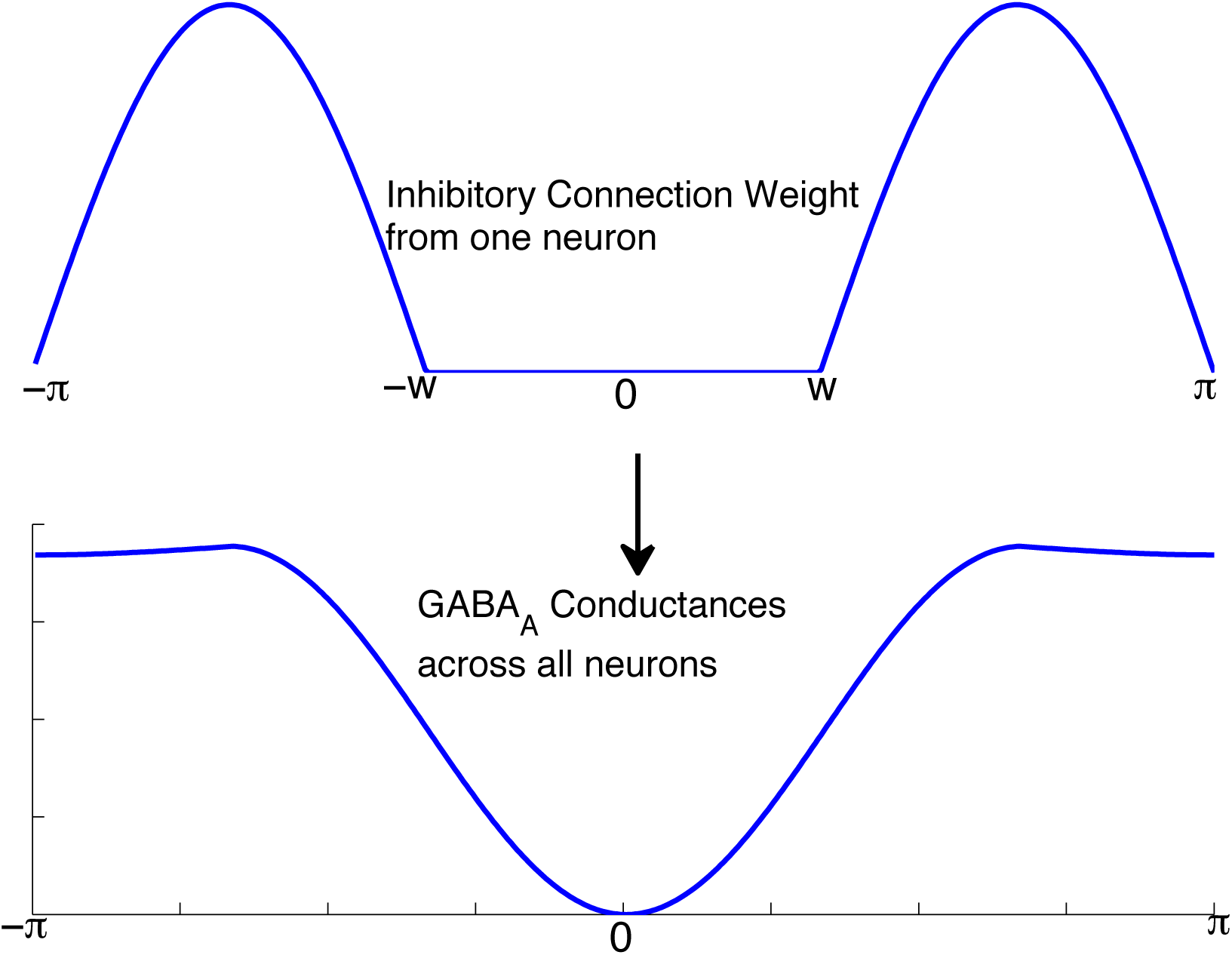
Definition for one dimensional cosine surround inhibition topology (top) based on the function *f*_*I*_ (*ϕ*) in the Appendix. It was constructed as two sine peaks with a gap 2*w* in between. Bottom curve is the *GABA*_*A*_ distribution across all neurons resulting from the surround inhibitory connectivities when the gap size parameter *w* = *π/*3 (see Figure 8). It is the result of the convolution of cosine surround function on the top with a spiking activity firing within the gap with a constant rate.

Convolution in Eq. (10) can be written down analytically and the resultant *g*_*GABA*_(*ϕ*) is a piece-wise function of *ϕ*. Below is the functional form of *g*_*GABA*_(*ϕ*) when *w ≤ π/*3 for *ϕ* = 0 to *π* (*g*_*GABA*_(*ϕ*) is symmetric with respect to *ϕ* = 0 so when *ϕ* = *−π* to 0 it is the mirror image of the following function):

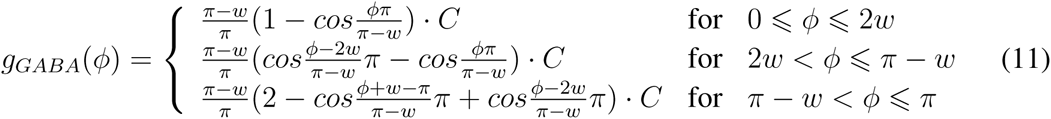

Analytic form of *g*_*GABA*_(*ϕ*) is plotted in Figure 7 as a surface plot for gap parameter *w* from 0 to *π/*3. We assumed the spiking bump activity occurs within a gap region centerred at *ϕ* = 0, as a result of surround inhibition, Figure 7 shows that neurons around that gap region around *ϕ* = 0 will have low GABA conductances which will allow the bump to emerge and potentially being stable. As described in the main text, other forms of inhibitory connectivity will not generate a local valley in the inhibitory conductances, thus destabilize any bump activity when it occurs. Figure 7 also shows that when the gap size 2*w* is small (e.g., *w < π/*6), neurons located at *π* and *−π* (they are equivalent because of wrap-around) will also have close to zero GABA inhibitory conductances. This suggests that a second bump could emerge and two bumps could coexist with half of the network size apart. Such phenomenon does occur in our simulations when using small gap size. In fact, in the two-dimensional neuronal array models, surround inhibitory topology is implemented as a taurus (we called it as Center-Annular-Surround (CAS) connectivity previously), two bumps normally emerge together with a distance apart. Figure 7 could potentially explain such fact because a second bump could emerge at half of the cycle (network size) apart because that is where inhibitory conductances could have another lowest value close to 0.

**Figure 8:**
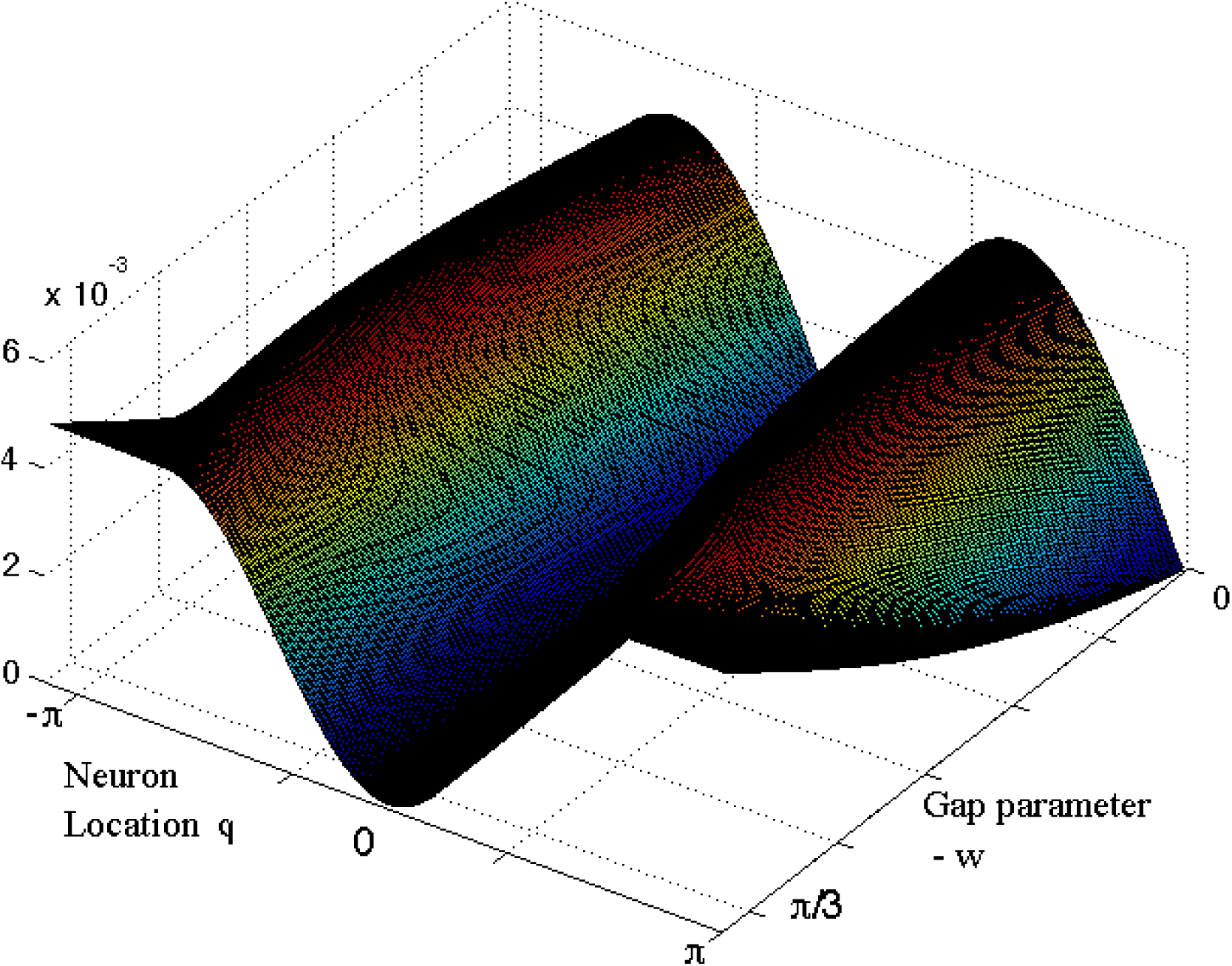
Normalized GABA conductance distribution function from Eq. (10) for gap size parameter *w* = 0 to *π/*3. The unit of the z-axis is normalized such that for every value of *w*, the sum of GABA conductance function from *−π* to *π* is normalized to 1. In actual numerical simulations this total sum of GABA conductance is a parameter (*S*_*I*_) on the x-axis in the transition curve on Figure 2 (d) to (f), and was scaled from 0 to 300 nS in our parameter scan for the transition curve.

As indicated in Eq. (4), in order for a bump to emerge, peak of the GABA conductances has to be larger than a certain value. This peak value can be found through Eq. (9). Firstly, we need to estimate the inhibitory firing rate within the bump area (*r*_*I*_ in Eq. (8)). Inhibitory firing rate *r*_*I*_ is determined by the currant input from the excitatory neurons (see Fig. 1), which in turn can be estimated by its excitatory NMDA conductance values. This is can be seen from Fig. 4(b), because of its longer time constant, NMDA conductances within the bump area continue increasing to a steady-state value after the bump is emerged. Because AMPA conductances have short time constant and decay back close to zero very fast after each spike from excitatory neurons, the steady-state values of the NMDA conductances for the inhibitory neurons will determine their firing rate. To estimate the inhibitory NMDA value, we assume firing rate of excitatory neurons within the bump area is *r*_*E*_ and it is related to excitatory AMPA and NMDA conductances through the conductance/firing rate curve (similar to the f/I curve): *r*_*E*_ = *f*_*fgcurve*_(*g*_*AMP A,E*_ + *g*_*NMDA,E*_). Fig. (9) summarizes the conductance/firing rate curve for the excitatory and inhibitory neurons we used in the network model analyzed here. From the figure we can see that above a certain conductance value, neuron’s firing rate roughly increases linearly with the increasing (excitatory) conductances for frequencies below 100*Hz*. This linear approximation can be used to estimate the firing frequency (such as *r*_*E*_ here) for a given AMPA and/or NMDA conductances.

From Eq. 6 we know the steady-state values of excitatory AMPA and NMDA conductances: *g*_*AMP A,E*_ +*g*_*NMDA,E*_ = *r_IN_ S_IN_ G_AMP A_τ*_*A*_ +*r_IN_ S_IN_ G_NMDA_τ*_*N*_. Based upon linear approximation of the firing frequency-conductances relations, excitatory firing rate will be proportional to the connection weights from the input layer:

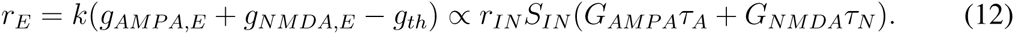

The most important message from the above equation is that firing rate of excitatory neurons in our network is not affected by the feedforward excitatory weight to inhibitory neurons (*S*_*E*_) nor the strength of the inhibitory weight (*S*_*I*_). This equation can explain the fact that excitatory firing rate in the WTA region within the *S*_*E*_ versus *S*_*I*_ parameter space in Figure 2(e) is mostly constant because we kept firing rate from the input layer and their connection weight the same while changing *S*_*E*_ and *S*_*I*_ systematically.

Since we have one-to-one feedforward connection from excitatory to inhibitory neurons, inhibitory NMDA conductances reach steady-state when the decay of NMDA conductances within an average period of input excitatory neuron (1*/r*_*E*_) is maintained by (and equal to) the amount of NMDA conductances generated by one excitatory spike, so suppose the peak value of inhibitory NMDA conductance is *x*, then:

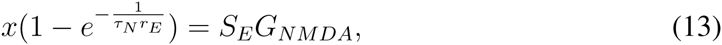

where *S*_*E*_ is the total feedforward excitatory connection weight to each inhibitory neuron, *G*_*N*_ is its associated NMDA gain factor representing the amount of NMDA conductance generated by one excitatory spike. The steady-state inhibitory NMDA conductances can be approximated by the decayed value of x:

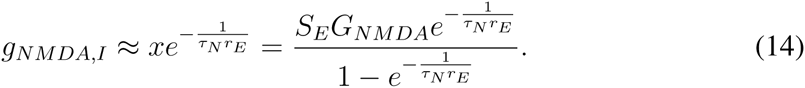

The inhibitory firing rate *r*_*I*_ then can be approximated by the neuron’s frequency-conductance curve relationship again:

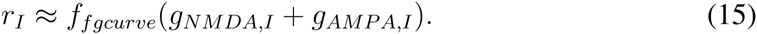

Based up Eq. (12), excitatory firing rate *r*_*E*_ is only related to firing rate of the input neurons, if we approximate the frequency-conductance curve by a linear relationship as shown in Fig. (9), and the steady-state value of AMPA conductances *g*_*AMP A,I*_ aprroximately cancel out the firing threshold *g*_*th*_, combining Eq. (12) and (13), inhibitory firing rate can be found to be proportional to the feedforward excitatory connection weight *S*_*E*_:

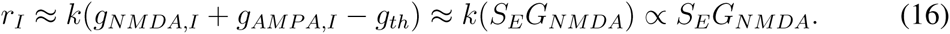

Above relation is confirmed by the observation in Fig. 2(f) that within the WTA parameter region, firing rate of inhibitory neurons increases as values of feedforward excitatory weight (*S*_*E*_ in the vertical axis) increases, while excitatory firing rate of WTA region (*r*_*E*_) in Fig. 2(e) remains close to a constant as indicated in Eq. (10).

Now using Eq. (9) we can estimate the maximal value of GABA conductances from the surround inhibition topology. If this maximal GABA value satisfy the condition on Eq. (4), then we can identify the locations of WTA region in the *S*_*E*_ versus *S*_*I*_ parameter space. For the gap size parameter we used (*w* = *π/*3), maximal GABA value can be approximated as the value of the function *g*_*GABA*_(*ϕ*) when *ϕ* = *±π* (see bottom trace in Fig. 8):

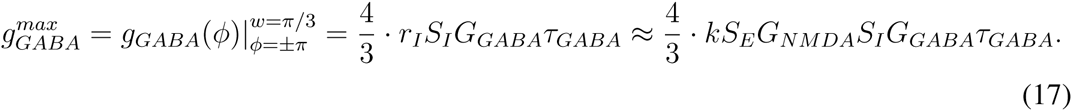

Combining results from the above equation and that from Eq. (4) which gives the threshold of peak GABA conductance value for a WTA bump to emerge, we now find the conditions for the parameters for the WTA region:

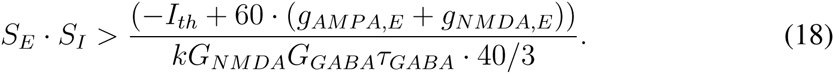

Most parameters on the right side of the above equation, i.e., *I*_*th*_ – firing threshold for input current, gain factors of NMDA and GABA conductances – (*G_NMDA_, G*_*GABA*_) and the slope for the conductance/firing rate curve (*k*), are all constants which can be derived or estimated from spike neuron model defined in Eq.(1). As we described before, AMPA and NMDA conductances for the excitatory neurons are only determined by the firing rate and the connection weight of the input neurons (see Eq. (8)). Using the relation in Eq. (8), the above transition condition can be expressed as the combination of the individual spiking neuron parameters and some information from the input layer:

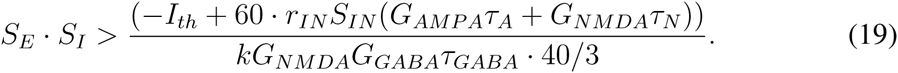

**Figure 9:**
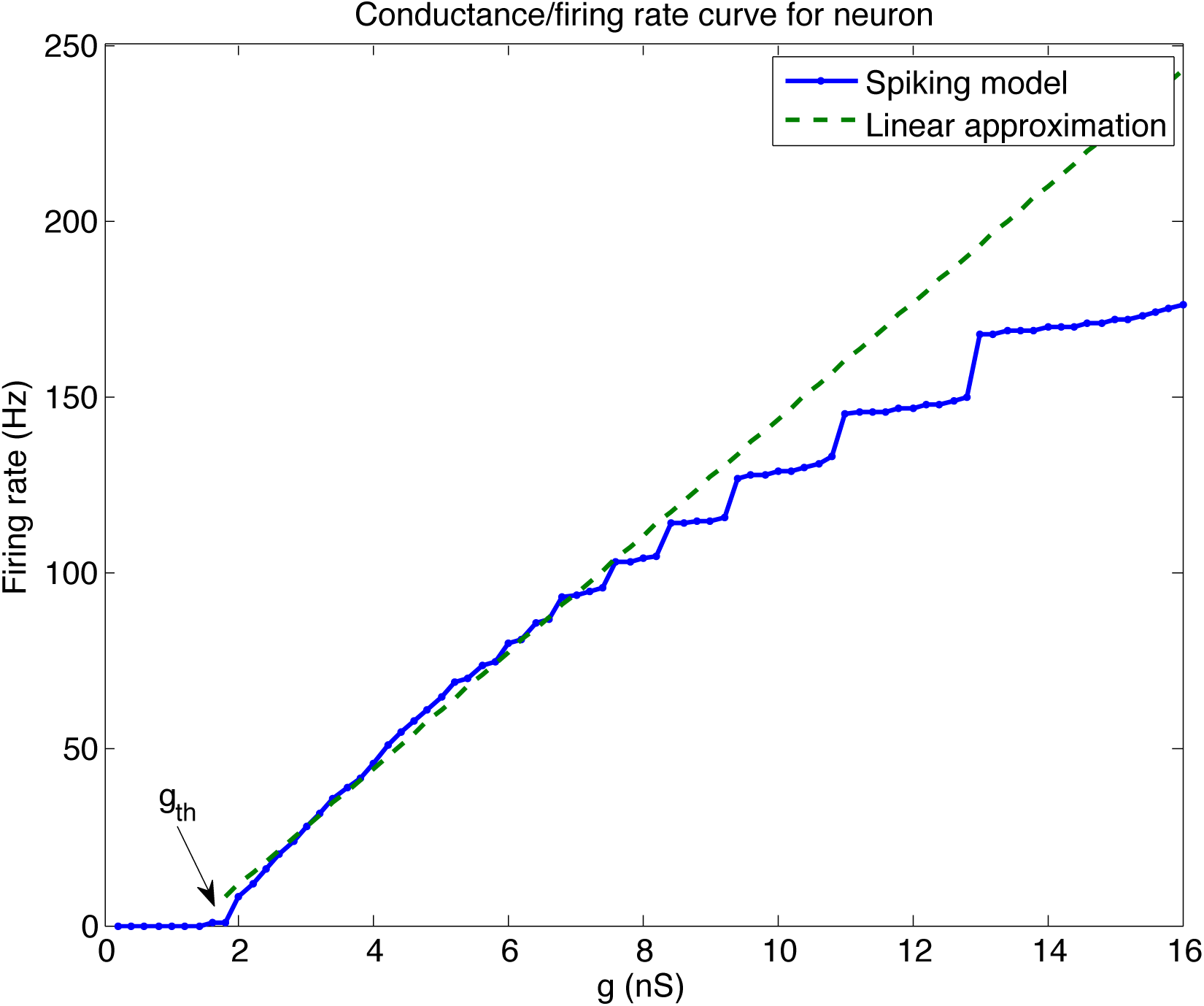
Firing rate versus (excitatory) conductance for the excitatory and inhibitory spiking neuron models we used in this paper. Above a certain threshold (*g*_*th*_), firing frequency roughly increases linearly with the increasing conductances, especially for the frequencies below 100Hz. Above that frequency, several plateaus appear and deviate from a simple linear relationship. Since neurons in our model mostly fire below 100Hz, which is the most biologically relevant frequency range, we can approximate this conductance/firing rate curve with a linear relation, *r* = *k*(*g − g*_*th*_) where *r* is a neuron’s firing rate, *k* is the slope, *g*_*th*_ is the threshold for firing. Such linear relation can be used in our analytical analysis to derive the transition curve for the Winner-Take-All state (see main text in the Appendix).

Eq. (19) thus reveals the analytic condition for the Winner-Take-All region in the parameter space, that is, multiplication of *S*_*E*_ (feedforward excitatory connection weight) and *S*_*I*_ (total sum of inhibitory connection weight) has to be larger than a constant value determined by individual neuron parameters and the input firing rate and connection weight. This equation suggests that the transition curve in the parameter space is an inverse function and gives the analytical explanations for the numerical results shown in the middle row of Fig.2. Based upon the specific parameters we used in out numerical simulation, *r_IN_ ≈* 14*Hz*, *S*_*IN*_ =100nS, *G*_*AMP A*_ = 0.15*nS/spike, G*_*NMDA*_ = 0.015*nS/spike, G*_*GABAA*_ = 0.15*nS/spike*, *τ*_*GABAA*_ = 6*msec.*, and the individual neuron parameters we estimated – *I*_*th*_ = 100*pA* and *k ≈* 37*Hz/nS*, we plotted the analytic condition for the WTA region using black curves in the middle row of the Fig. 2 (– (d), (e), (f)). As we can see, our analytic solution fits the numerical results very well and clearly locates the winner-take-all region in the parameter space.

Above Eq. (19) is obtained under the condition of considering only one GABA conductance (i.e., *GABA*_*A*_). If we include both *GABA*_*A*_ and *GABA*_*B*_ conductances such as defined in Eq. (2), we can derive the conditions for the WTA state including all four type of conductances using relations in Eq. (5):

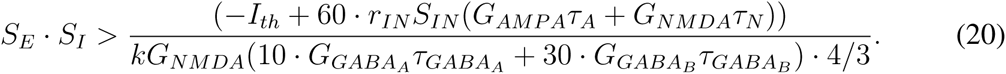

By adding *GABA*_*B*_ conductances and increasing the denominator in the above equation we can see that the transition curve defined by Eq. (20) will be shifted towards bottom-left in the *S*_*E*_ versus *S*_*I*_ parameter space and the winner-take-all region will be increased. This is indeed the case. Fig. (10)(a) show the maximal firing rate of excitatory neurons and the WTA region when both *GABA*_*A*_ and *GABA*_*B*_ conductances are included in our model. White dashed line is based upon condition specified in Eq. (20). Compared with Fig 2(d), we can see the WTA region has greatly increased and the transition curve has been shifted closer to the origin, so much that the WTA region on the bottom-right side has been merged with the previous separated and disconnected high-firing region. However, an interesting observation is that there are three or more blue regions (low firing rate) beyond the white transition curve calculated based upon Eq. (20). Closer inspection reveals that these are actually stable epileptic states (synchronized rhythmic firing between excitatory and inhibitory neurons) which are previously outside of the WTA region in Fig 2(d). More interestingly, three different regions correspond to different ratios of firing rates between excitatory and inhibitory neurons (i.e., 1:1 rhythmic in Fig. (10)(b), 1:2 rhythmic in Fig. (10)(c) and 1:3 rhythmic in Fig. (10)(d) respectively). In fact, if we scan the parameter space in higher resolutions, we might locate (blue) regions with higher frequency ratios (e.g., 1:4). We believe what we observed here is the “Arnold Tongue” region (Coombes & Bressloff, 1999) for this spiking network.

**Figure 10:**
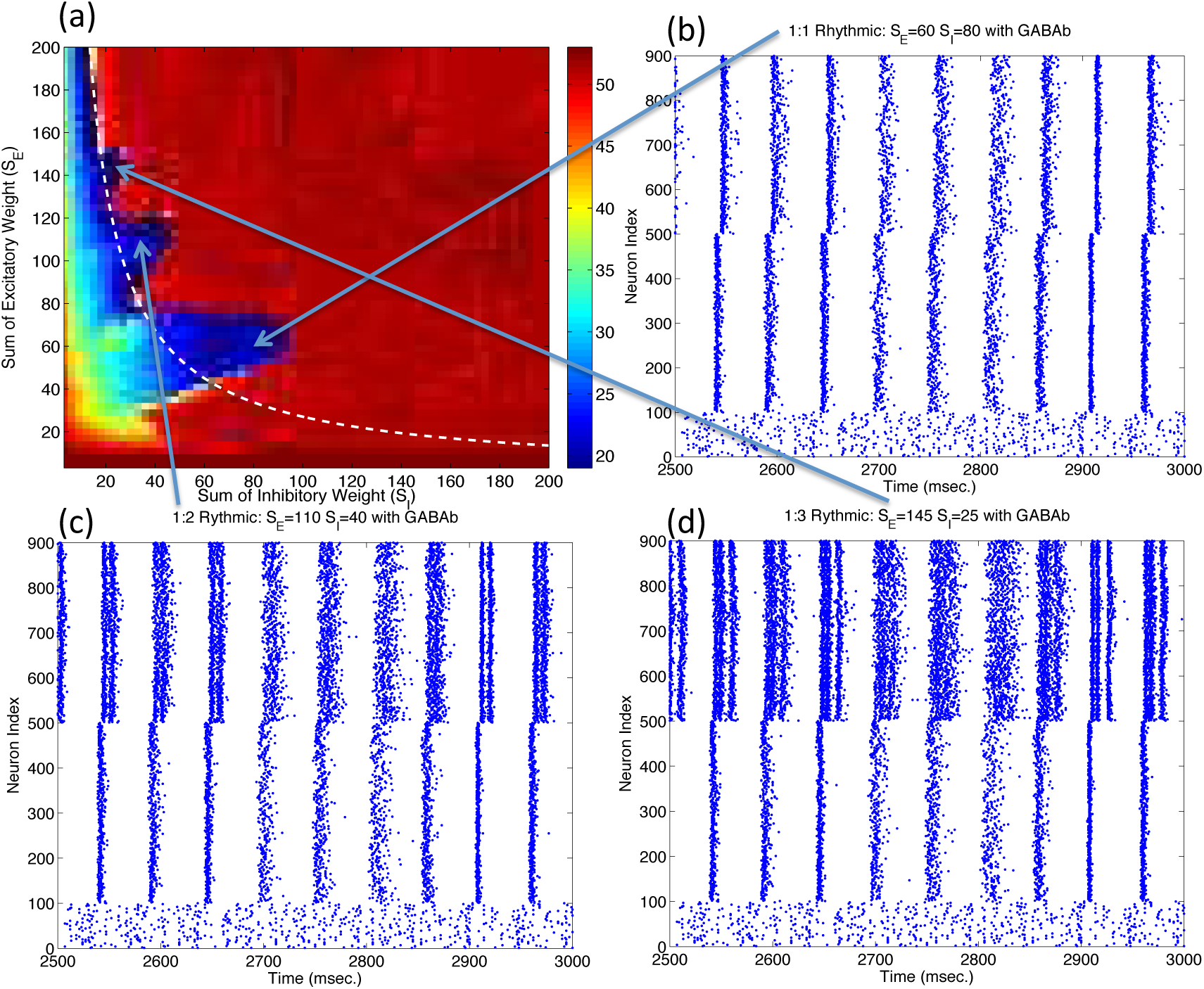
Arnold tongue and the increased Winner-Take-All region in the parameter space with both *GABA*_*A*_ and *GABA*_*B*_ conductances.

